# Tectal stem cells display diverse regenerative capacities

**DOI:** 10.1101/268136

**Authors:** Benjamin W. Lindsey, Georgia E. Aitken, Jean K. Tang, Mitra Khabooshan, Celia Vandestadt, Jan Kaslin

## Abstract

How diverse adult stem and progenitor populations regenerate tissue following damage to the CNS remains unknown across most neurogenic domains. To understand the role of quiescent radial-glial (qRG) stem cells during regeneration, we tested the hypothesis that qRG could be induced to proliferate and produce newborn neurons. We designed a stab lesion assay in the midbrain tectum of the adult zebrafish to target an isolated population of qRG, and investigated their proliferative behaviour, differentiation potential, and requirement of Wnt/β-catenin signalling for the regenerative response. EdU-labelling showed that a small proportion of qRG transit to a proliferative state (pRG), but that progeny of pRG are restricted to a radial-glial fate. Lesion promoted upregulation of proliferation and neurogenesis from neuro-epithelial-like amplifying progenitors (NE-Ap) of the tectal marginal zone (TMZ). Homeostatic levels of Wnt/β-catenin signalling persisted under lesioned conditions in the qRG/pRG population, whereby increased β-catenin staining and *axin2* expression was present in the NE-Ap progenitor zone. Attenuation of Wnt signalling using *Dickkopf-1*, demonstrated that proliferative responses post-injury appeared to be Wnt-independent. Our results align with the emerging view that adult stem/progenitor phenotypes are characterized by discrete, rather than mutual, regenerative programs and that different stem cell domains employ different modes of regeneration.

## INTRODUCTION

The adult stem cell niche is composed of heterogeneous neural stem and progenitor cells, reflecting their developmental origin, cell lineages, and proliferative dynamics (Lindsey and Tropepe, 2006; Grandel et al., 2006; Adolf et al., 2006; Shen et al., 2006; Kaslin et al., 2009; Kriegstein and Alvarez-Buylla, 2009; Grandel and Brand, 2013; Giachino et al., 2014; Furutachi et al., 2015; Than-Trong and Bally-Cuif., 2015; Chaker et al., 2016). Presently, the cellular and molecular signatures of these populations are best understood under homeostasis and repair within the vertebrate forebrain niche (Merkle et al., 2007; Marz et al., 2010; 2011; Ganz et al., 2010; 2012; Rothenaigner et al., 2011; Kroehne et al., 2011; Ihrie and Alvarez-Buylla, 2011; Kishimoto et al., 2011; 2012; Lindsey et al., 2012; Codega et al., 2014; Fuentealba et al., 2015; Luo et al., 2015; Llorens-Bobadilla et al., 2015). Outside of the forebrain we are only beginning to uncover the regenerative plasticity of stem/progenitor cells and their biological importance (Magnusson and Frisén, 2016), particularly in highly regenerative vertebrates.

The zebrafish has emerged as a leading model of stem cell plasticity and regeneration with its wealth of neurogenic compartments positioned along brain ventricles (Zupanc et al., 2005; Kaslin et al., 2008; Ito et al., 2010; Kizil et al., 2012; Lindsey and Tropepe, 2014; Lindsey et al., 2015; 2018; Alunni and Bally-Cuif, 2016; Ghosh and Hui, 2016). Niches are populated by heterogeneous stem/progenitor phenotypes, dominated largely by neuro-epithelial-like (NE) stem/progenitor cells and radial-glial cells residing in proliferative (pRG) or quiescent (qRG) states. The striking regenerative ability of the zebrafish brain has given rise to the notion that most adult stem/progenitor cells are likely to be multipotent, and as such, capable of replacing all cell lineages lost during injury (i.e. NE, RG, oligodendrocytes, neurons). While this hypothesis appears to be upheld by quiescent Müller glia of the adult retina (Otteson and Hitchcock, 2003; Ramachandran et al., 2011; Wan et al., 2015), the unique regenerative profile of individual cell phenotypes across distinct stem cell niches of the brain is less clear.

Radial-glia of the dorsal telencephalon have been the focus of most injury studies in the zebrafish CNS (Marz et al., 2011; Kroehne et al., 2011; Kyritsis et al., 2012; Baumgart et al., 2012; Kishimoto et al., 2012, Barbosa et al., 2015; reviewed in Alunni and Bally-Cuif, 2016; Lindsey et al., 2018). We have shown that RG play a major role in regenerating new neurons that repopulate those lost, with these cells fated to become functional neuronal subtypes (Kroehne et al., 2011). Interestingly, under homeostasis a large proportion of the dorsal RG population remain quiescent, regulated by strong expression of Notch genes (Chapouton et al., 2010; Rothenaigner et al., 2011). Downregulation of Notch signalling induces qRG to re-enter the cell cycle and increase symmetric divisions (Alunni et al., 2013), allowing them to respond to injury. In contrast, within the cerebellar niche RG are quiescent and do not serve as functional stem cells, with neurogenesis driven solely by multipotent NE-like stem cells under homeostasis (Kaslin et al., 2009). We recently revealed that upon surgical injury to the cerebellum, tissue regeneration was governed primarily by the NE-like population despite re-entry of qRG into the cell-cycle, recapitulating the homeostatic state of the niche (Kaslin et al., 2017).

Distinct from other major structures of the adult CNS, the zebrafish midbrain tectum contains stem cell niches populated entirely by a single stem/progenitor cell type. Here, an extensive population of qRG exist at the roof of the tectal ventricle, while neuro-epithelial-like amplifying progenitors (NE-Ap) that contribute to lifelong neurogenesis are found at the tectal marginal zone (TMZ; Ito et al., 2010). Embryonically these cells are derived from slow-amplifying progenitors (Recher et al., 2013). Recently it has been demonstrated that these NE-Ap are the final progenitor phenotype of a well-defined NE lineage that originate from *Her5-*positive NE stem cells in the posterior tectum (Galant et al., 2016). Moreover, recent evidence suggests that NE cells of the zebrafish tectum rely on Wnt/β-catenin signalling to promote stem cell proliferation towards a neuronal phenotype (Shitasako et al., 2017), similar to the role of Wnt in the adult mammalian stem cell niches (Adachi et al., 2007; Jang et al., 2013; Seib et al., 2013). At present, the regenerative capacity of qRG and NE cells within the midbrain tectum (**Fig. 1c**), along with the signalling pathways governing their regulation, remain largely unknown.

**Figure 1.**
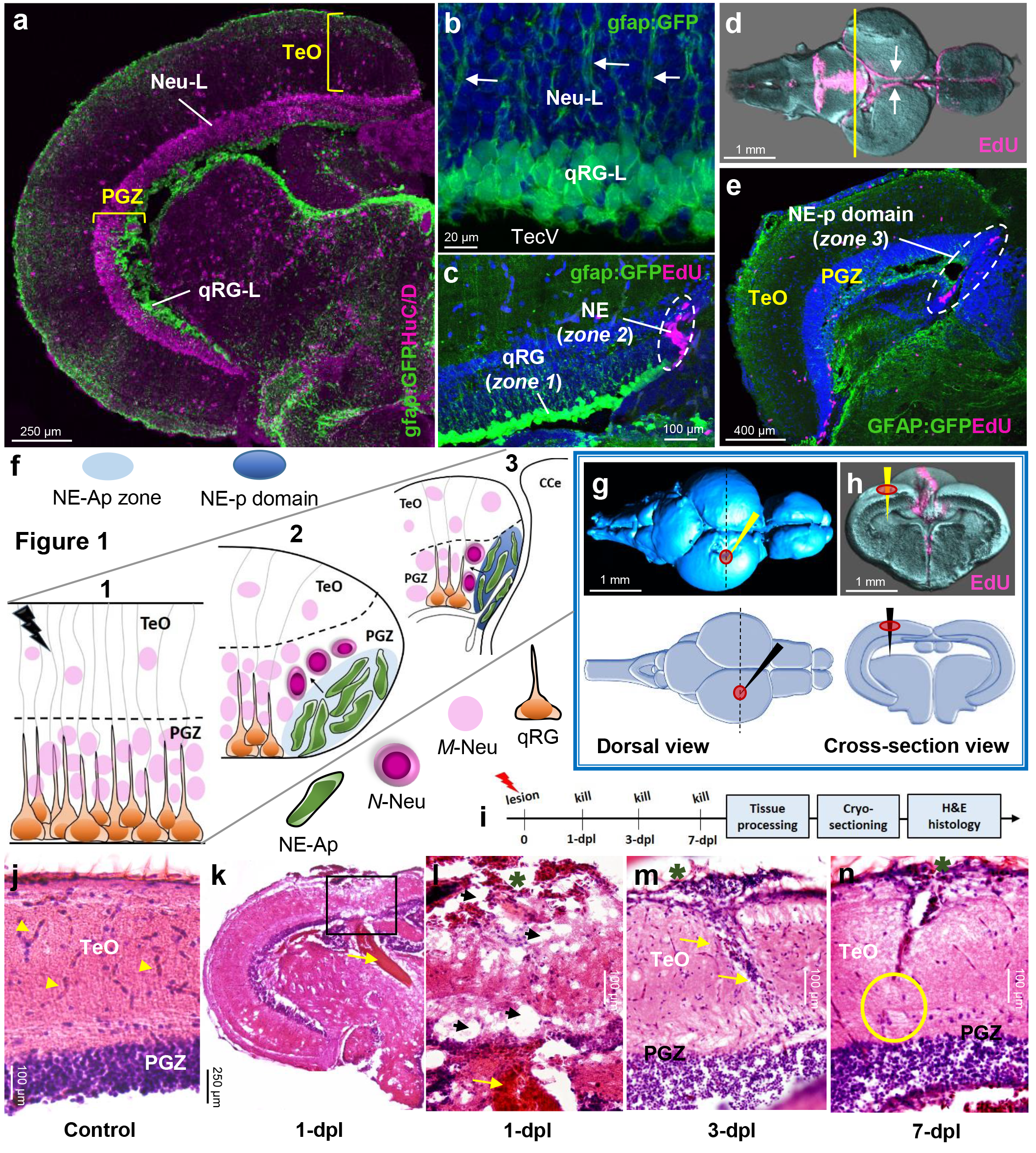
Tectal composition, stem cell niches, and stab lesion model. **(a)** Cross-sectional view of a single adult tectal hemisphere showing a laminar composition consisting of outer tectal superficial layers (TeO) and the deeper, cell-dense, periventricular grey zone (PGZ). The PGZ is subdivided into an upper neuronal layer (Neu-L, HuC/D labelling; pink) and a quiescent radial-glial layer (qRG-L; *Tg*(*gfap:GFP*)*^mir2001^* labelling; green) populating the roof of the tectal ventricle. **(b)** High magnification of the PGZ illustrating the 3-4 cell deep structure of the qRG-L (green) abutting the tectal ventricle (TecV) with radial processes extending upwards from qRG cells through the Neu-L (white arrows) and towards the superficial tectal layers. **(c)** Neuro-epithelial-like (NE; pink; zone 2) stem/progenitor cells identified by EdU-labelling are located at the tectal marginal zone (TMZ; white dashed circle), adjacent to the qRG population (green; zone 1). **(d)** Dorsal view of the homeostatic staining pattern of cell proliferation using EdU (pink) in reconstructed whole brains following optical projection tomography, showing constitutively proliferating NE stem/progenitor cells along the length of the tectal margin (white arrows).Yellow line depicts cross-sectional level shown in panel **(e)**. **(e)** Cross-section view of the NE progenitor domain (NE-p; white dashed circle; zone 3) labelled with EdU (pink), from where the NE-lineage and RG-lineage arise, residing at the posterior aspect of the adult midbrain adjacent to the medially located cerebellum. **(f)** Schematic representing the three stem cell zones (1-3) investigated following tectal stab lesion (lightning bolt) superficial to the underlying qRG layer: 1 – qRG population lining the tectal ventricle; 2 – constitutively proliferating NE amplifying progenitor zone (NE-Ap zone; light blue) located at the tectal marginal zone (TMZ) adjacent the qRG population; 3 – posterior NE-p domain (dark blue). Note that continuously proliferating NE cells in zones 2-3 produce newborn neurons (*N*-Neu) under physiological conditions. *M*-Neu, mature neuron; CCe, corpus cerebelli **g-h:** Dorsal (**g**) and cross-sectional (**h**) views of the intact control brain depicting the neuroanatomical level (**h**, black dashed lines) and cannula insertion site (red circles/black or yellow arrows) for our tectal stab lesion into the left hemisphere in OPT reconstructed brains (top) and schematic images (bottom). **I:** Experimental design for hematoxylin and eosin (H&E) staining following lesion. **J:** Control tectum showing well organized PGZ and highly vascularized superficial TeO layers (yellow arrowheads). **k-l:** Tissue 1-dpl showing extensive damage across tectal layers (TeO + PGZ) following cannula insertion, presence of vacuoles at the lesion site, and extensive blood pooling underlying the fragmented PGZ (yellow arrows). Black box depicts higher magnification shown in **(k)**. **(m)** Tissue at 3-dpl was characterized by a reduction in the size of vacuoles and oedema, along with a pronounced increase in the number of cell bodies within the lesion canal (yellow arrows). **(n)** Tissue at 7-dpl demonstrated a reduction in the number of cell bodies from within the lesioned canal, with repair of the canal occurring first at the bottom half of the superficial tectal layers (TeO; yellow circle). Surrounding tissue resembles more closely the uninjured state. In panels **m-n** green asterisks denote the location of lesion. Cross-sectional views are shown for all histological images.

In this study, we designed a tectal stab lesion to examine the proliferative dynamics, neurogenic potential, and requirement of Wnt/β-catenin signalling during the regenerative response in three distinct stem cell zones (**Fig. 1f**): populations of qRG (zone 1), NE-Ap (zone 2), and NE progenitor (NE-p) in the posterior tectal domain (zone 3). We show that while a modest population of normally quiescent ventricular RG can be stimulated to re-enter the cell cycle and produce EdU^+^ progenitors, these progenitors are restricted to producing RG. No new neurons were detected at the lesion site after injury. However, mechanical injury was sufficient to promote upregulation of constitutive levels of proliferation and neurogenesis from NE-Ap at the TMZ, implicating this population in neuroregeneration as previously demonstrated in the cerebellum. Downstream targets of Wnt/β-catenin signalling, including β-catenin and *axin2*, remained unchanged between uninjured and injured states in the qRG layer of the midbrain, but were elevated in the NE-Ap zone. Following tectal lesion attenuation of Wnt/β-catenin signalling via its antagonist, *Dickkopf-1*, revealed no change in the proliferative response of EdU^+^ cells in the quiescent RG layer of the PGZ, nor the NE-Ap population at the TMZ, suggesting a Wnt-independent proliferative response. Taken together, our results contribute to the notion that the regenerative potential of the adult zebrafish CNS is orchestrated by discrete stem/progenitor phenotypes that are regulated by intrinsically-defined regenerative programs.

## RESULTS

### The adult optic tectum contains an isolated population of non-cycling quiescent radial-glial cells lining the roof of the tectal ventricle

Quiescent radial-glial (qRG) stem/progenitor cells comprise a proportion of many heterogeneous adult stem cell niches throughout the teleost brain (Kaslin et al., 2009; Alunni et al., 2010; Chapouton et al., 2010; Lindsey et al., 2014). Within the optic tectum, an extensive, isolated population of qRG cells are uniquely positioned adjacent to the tectal ventricle. These constitute the quiescent radial-glial layer (qRG-L) below the upper neuronal layer (Neu-L) of the periventricular grey zone (PGZ; **Fig. 1a**). Reporter lines for glial fibrillary acidic protein (GFAP) and *her4.1*, as well as antibodies against glial markers glutamine synthetase (GS), S100β, and fabp7a (Ito et al., 2010), show that the qRG-L is 2-3 cells deep. Individual cell processes of these qRG project upwards through the multiple layers of the tectum to the superficial tectal layer (Stevenson and Yoon, 1982; Bartheld and Meyer, 1987; **Fig. 1b**, white arrows). Injections with the *S*-phase cell cycle marker, 5-ethynyl-2`-deoxyuridine (EdU), demonstrate that unlike actively proliferating RG cells (pRG), glia of this layer do not exist within a proliferative state (**Fig. 1c**; qRG; Ito et al., 2010). EdU, however, is incorporated by neuroepithelial-like (NE) progenitor cells located at the tectal marginal zone (TMZ) which contribute to constitutive neurogenesis throughout life (Ito et al., 2010; **Fig. 1c**, NE). Whole brain EdU-staining shows that NE amplifying progenitors (NE-Ap) are situated along the length of the adult TMZ, forming a ribbon of cells at the medial edge of each hemisphere that continues posteriorly, adjacent to the cerebellum (**Fig. 1d**, white arrows). Cross-sections through the posterior tectum reveal that at this neuroanatomical level (yellow line), a larger domain of EdU^+^ NE progenitor cells can be identified (**Fig. 1e**; dashed lines), from which the tectal NE lineage and RG cells are derived (Galant et al., 2016). These distinctive stem/progenitor niches within the adult optic tectum provide an exceptional experimental system to investigate how each population (**Fig. 1f**; ***1 – qRG; 2-NE-Ap, 3-NE-p domain cells;* see also** **Fig. 1c, e** for location of zones) responds to injury and contributes to regenerative neurogenesis.

To address this question, we designed a tectal stab lesion assay whereby a cannula, positioned dorsomedially in a single tectal hemisphere was inserted vertically into the neurocranium through the tectal layers, terminating at the ventricle (**Fig. 1g-h**; top: OPT-rendered brain; bottom: schematic view of lesion coordinates). We reasoned that the location of our injury site was optimal to elicit a response from the underlying qRG cells in the PGZ, and was in close enough proximity to the NE-Ap population for these cells to respond to injury-induced signals. Moreover, the posterior directional flow of cerebrospinal fluid through the tectal ventricle provides a direct path of transmission of signals arising from the lesion site to contact NE cells adjacent to the ventricle in the posterior NE-p domain. Three-dimensional sphere analysis of the distribution of EdU^+^ staining radiating from the lesion site over the first 3-days post-injury (**Fig. S1j;** control, *n* = 6 brains; 1-dpl, *n* = 10 brains; 3-dpl, *n* = 8 brains) revealed a progressive decline in the proportion of EdU volume/sphere at 1-dpl from ~ 45 % to 25%, that was sustained thereafter (**Fig. S1k-l**). By contrast, we detected a significant reduction in the proportion of EdU volume/sphere between the sphere nearest the lesion site (i.e. 0-49 μm sphere) and those more distal at 3-dpl (**Fig. S1m-n**), suggesting that injury-induced cell proliferation was most pronounced nearest the lesion site at all time points. Comparison across time points further illustrates that the proliferative response to tectal injury is markedly higher distal to the lesion site at 1-dpl (*see* **Fig. S1k**). Antibody labelling at 3-dpl using Proliferating Cell Nuclear Antigen (PCNA), a cell cycle marker of G1 and subsequent phases, showed that the PGZ at cross-sectional levels anterior and posterior to the lesion site (~ 50 μm +/− lesion site) had pronounced levels of PCNA-positive cells in the lesioned hemisphere (data not shown). This demonstrates that signals arising from the lesion site produce a far-reaching proliferative response. These findings led us to ask whether the global proliferative response of the brain to injury might be conserved independent of the lesion site. By comparing our telencephalic (Kroehne et al., 2011) and tectal stab lesion assays following OPT whole brain staining and EdU imaging (Lindsey et al., 2018; Lindsey and Kaslin, 2017), we consistently observed upregulation of EdU at 1-dpl across the neuro-axis before becoming restricted to the lesion site at 3-dpl, and resembling uninjured (i.e. control) levels by 7-dpl (**Fig. S2a-b**). This result may imply that the vertebrate brain activates an acute, brain-wide stereotypical proliferative response to injury, prior to acting specifically at the site of injury.

Histological analysis of the tissue lesion over the first week post-injury using hematoxylin and eosin (H&E) staining (**Fig. 1i**; control, *n* = 3; 1-dpl, *n* = 5; 3-dpl, *n* = 5; 7-dpl, *n* = 4; 3-mth, *n* = 3) demonstrated that compared with well delineated tectal layers in control tissue (**Fig. 1j**), a conspicuous stab canal along with disorganised tectal layers was present by 1-dpl (**Fig. 1k**). The injured hemisphere appeared highly disrupted at the lesion site with extensive bleeding leading to the presence of oedema extending ventrally from the tectal ventricle into the underlying parenchyma (**Fig. 1k-l**; yellow arrows). Compared with control (*see* **Fig. 1j**) and uninjured tissue adjacent the lesion site at 1-dpl, we further detected an increase in the number of vacuoles (black arrowheads) and a loss of vascularization (**Fig. 1l**). At 3-dpl blood clotting was no longer present, damaged tissue was less vacuolated, and the vertical lesion formed by the cannula clearly visible and filled with a conspicuous number of cell bodies (**Fig. 1m**; yellow arrows). This prompted us to ask whether the extensive number of cells within the lesion site might be a result of the rapid immune response known to occur following CNS injury. Indeed, examination of the tectal injury site using the *Tg*(*mpeg1:mCherry*)^*g122*^ transgenic reporter line that labels brain microglia and peripheral macrophages (**Fig. S3a**; Ellett et al., 2011; control, *n* = 3; 1-dpl, *n* = 4; 3-dpl, *n* = 4), showed that unlike control conditions where few resident microglia were present (**Fig. S3b**), at 1-dpl a large number of macrophages were recruited to the lesion site and lined the vertical canal (**Fig. S3c**). At 3-dpl the majority of these cells were located within the vertical canal (**Fig. S3d**), accounting, at least in part, for the increase in cell bodies observed from histological analysis at this same time point. Along with recruitment of macrophages towards the lesion canal following injury we also noted a change in the cellular morphology of macrophages from enlarged cell bodies with ramified processes (***see*** **Fig. S3b;** red circle) to a condensed, ameboid-like morphology (*see* **Fig. S3c**; red circle; Svahn et al., 2013). This is indicative of a shift from a surveillance role under homeostasis to activation of these macrophages post-lesion. EdU labelling demonstrated that compared with unlesioned tissue, where microglia remain non-proliferative (**Fig. S3e**), a small fraction of macrophages (microglia or infiltrated peripheral macrophages) appeared to be double-labelled with the proliferative marker at 1-dpl and 3-dpl (**Fig. S3f-g**; white arrows). By 7-dpl, our histological analysis displayed a reduction in the size of the lesion canal in the more inferior tectum (yellow circle) along with a notable decrease in eosin-positive cell bodies (**Fig. 2n**). By 3-mth post-injury tissue from both sides of the lesion canal had bridged together, with tectal layers beginning to more closely resemble their stereotypical laminar organization (data not shown).

**Figure 2.**
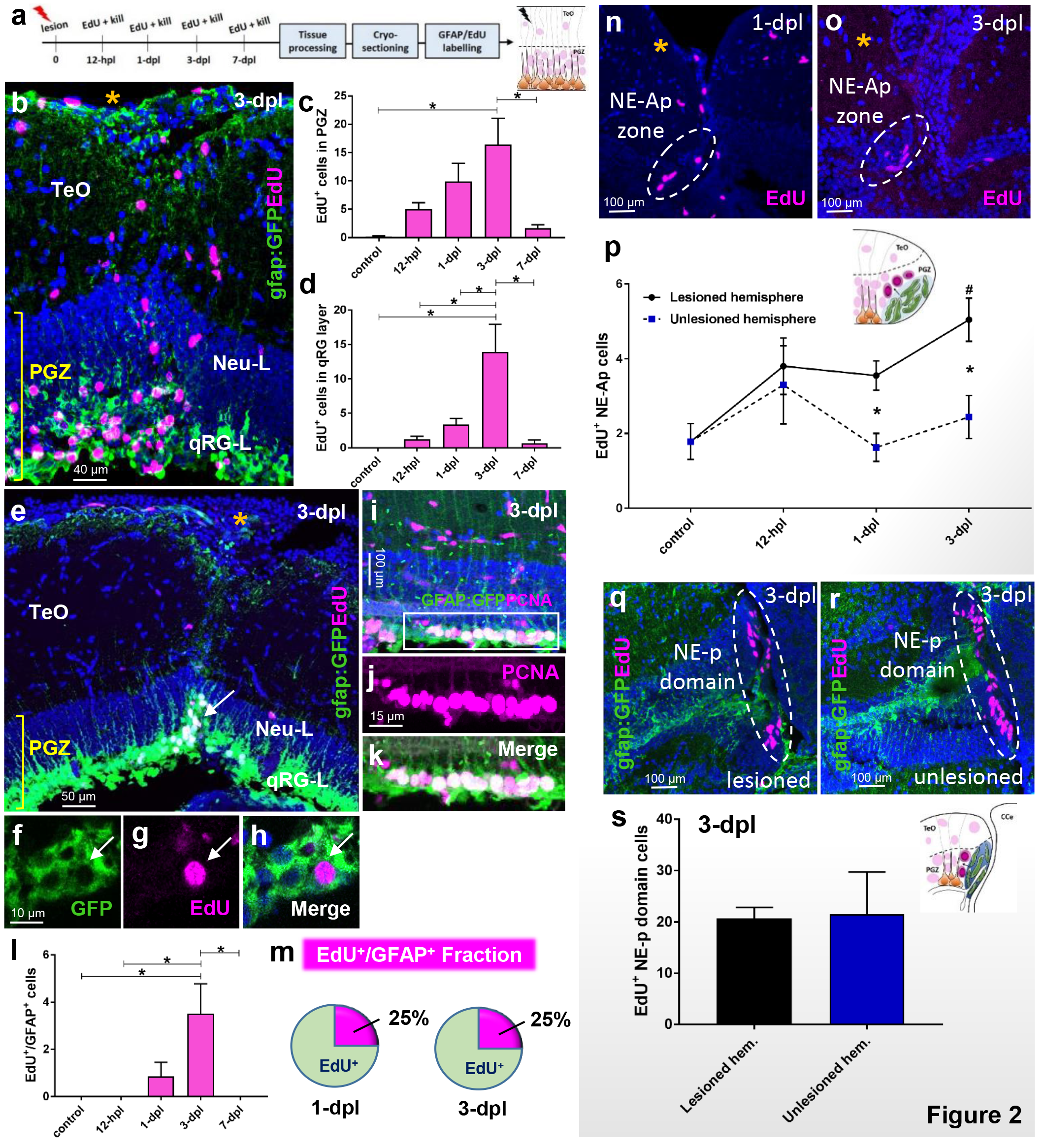
Cell proliferation post-lesion in stem cell zones. **(a)** Experimental design to investigate EdU proliferation arising from the qRG and NE populations. **(b)** Example of the proliferative response at the lesion site (orange asterisk) in the PGZ and superficial layers at 3-dpl when response is maximal. **(c)** Total EdU^+^ cells in the PGZ (Neu-L + qRG-L) at times post-lesion. One-way ANOVA; F (4, 23) = 4.395, *p* = 0.0087; Tukey’s multiple comparisons test: 3-dpl vs control, *p* = 0.0109; 3-dpl vs 7-dpl, *p* = 0.0365. **(d)** Total EdU^+^ cells in the qRG layer at times post-lesion. One-way ANOVA; F (4, 24) = 8.585, *p* = 0.0002; Tukey’s multiple comparisons test: 3-dpl vs control, *p* = 0.0005; 3-dpl vs 12-hpl, *p* = 0.0008; 3-dpl vs. 1-dpl, *p* = 0.0031; 3-dpl vs 7-dpl, *p* = 0.0018. **(e)** Example of a population of co-labelled GFAP:GFP^+^/EdU^+^ cells in the qRG layer of the PGZ at 3-dpl (qRG-L, white arrow). High magnification view of proliferating radial-glia (pRG) in separate GFP (**f**) and EdU (**g**) channels, and merge (**h**) at 3-dpl. **(i-k)** Co-labelled population of GFAP:GFP^+^/PCNA^+^ pRG lining the tectal ventricle at 3-dpl. White box in **i** denotes images shown in **j** and **k**. **(l)** Total EdU^+^/GFAP^+^ proliferating radial-glial (pRG) cells arising from activated qRG at times post-lesion. One-way ANOVA; F (4, 22) = 4.479, *p* = 0.0085; Tukey’s multiple comparisons test: 3-dpl vs control, *p* = 0.0308; 3-dpl vs 12-hpl, *p* = 0.0133; 3-dpl vs. 1-dpl, *p* = 0.0707; 3-dpl vs 7-dpl, *p* = 0.0308. **(m)** Fraction of EdU^+^/GFAP^+^ cells as a percentage of the total EdU^+^ population in the PGZ at 1-dpl and 3-dpl. **(n-o)** Examples of the EdU^+^ population size of NE-Ap cells (dashed white circles; pink EdU) observed in the lesioned (yellow asterisk) and unlesioned hemispheres at 1-dpl (**n**), and 3-dpl (**o**). **(p)** Number of EdU^+^ NE-Ap cells in lesioned hemispheres at times post-injury compared with control levels (one-way ANOVA, F (3, 18) = 7.686, *p* = 0.0016; Tukey’s multiple comparisons test, *p* = 0.005), and between lesioned and unlesioned hemispheres at each time point (unpaired t-test, two-tailed: 1-dpl lesioned vs 1-dpl unlesioned, p = 0.0061; 3-dpl lesioned vs 3-dpl unlesioned, p = 0.0096). **(q-r)** Examples of EdU^+^ labelling (pink) in the NE-p domain of the posterior tectum (white dashed circles) at 3-dpl between the lesioned and unlesioned hemispheres. **(s)** Quantification of the number of EdU^+^ cells between the lesioned and unlesioned hemispheres of the posterior NE-p domain (unpaired t-test, two-tailed: *p* = 0.9211). All data presented are mean ± S.E.M. Significance was accepted at \**p* < 0.05.

### Injury stimulates a subpopulation of quiescent radial-glial cells to re-enter the cell cycle and upregulates proliferation from neuroepithelial-like amplifying progenitors

How populations of qRG and progenitors along the NE lineage modulate their proliferative behaviour upon lesion remains poorly understood. To examine the proliferative response of qRG and NE progenitor cells following lesion, we performed EdU-labelling in the *Tg(gfap:GFP*)*^mir2001^* reporter line between 12-hours post lesion (hpl) and 7-dpl (**Fig. 2a**; control, *n* = 5; 12-hpl, *n* = 5; 1-dpl, *n* = 8; 3-dpl, *n* = 5; 7-dpl, *n* = 4). Using 4-hr EdU-chase periods we observed a progressive increase in the proliferative response in the PGZ (qRG-L + Neu-L) that peaked at 3-dpl before significantly decreasing at 7-dpl (**Fig. 2b-c**). The same trend in EdU-labelling as observed in the entire PGZ was seen within the quiescent RG layer (qRG-L) of the tectum, with a significant increase in the EdU population size at 3-dpl compared with all other groups (**Fig. 2d**). Importantly, from the qRG layer we successfully identified a subset of constitutively quiescent RG that newly entered the cell cycle and were EdU^+^/gfap:GFP^+^ (**Fig. 2e-h**). We further confirmed this finding using PCNA, illustrating an even greater number of newly proliferative co-labelled RG cells (**Fig. 2i-k**). Analysis of the EdU^+^/gfap:GFP^+^ population showed that these proliferating RG, hereafter referred to as “pRG”, were restricted to a 48-hr window spanning 1-dpl through to 3-dpl. The presence of pRG were maximal at 3-dpl and significantly different from most other groups (**Fig. 2l**). Interestingly, despite an approximate doubling of the number of EdU^+^/gfap:GFP^+^ cells from 1-dpl to 3-dpl, the fraction of EdU^+^/gfap:GFP^+^ cells compared to the total EdU population remained steady at 25% (**Figure 2m**). These experiments highlight the ability of a subpopulation of qRG to be stimulated to enter a proliferative state as a result of physical injury.

Analysis of NE-Ap at the tectal margin (control, *n* = 8; 12-hpl, *n* = 3; 1-dpl, *n* = 5; 3-dpl, *n* = 6) and cells within the posterior NE-p domain (3-dpl, *n* = 4) showed that tectal injury had differential effects on the proliferative behaviour of these two populations. We detected a significant increase in EdU^+^ cells at 1-dpl and 3-dpl between lesioned and unlesioned hemispheres in NE-Ap (**Fig. 2n-p**; dashed line), in addition to a significant increase between homeostatic levels of proliferation (i.e. control) from NE-Ap and the lesioned hemisphere at 3-dpl (**Fig. 2p**). Conversely, levels of proliferation remained unchanged in the more posterior NE-p domain between lesioned and unlesioned hemispheres when quantified at 3-dpl (**Fig. 2q-s**; 3-dpl, *n* = 4). These results imply that injury acts specifically on NE-Ap cells proximal the lesion site, rather than the mixed NE lineages that make up the posterior NE-p domain (Galant et al., 2016). Given these findings, we specifically targeted only the qRG population and NE-Ap zone in subsequent experiments.

### Progeny of proliferative radial-glial produce only newborn glia, whereby neurogenic output from neuroepithelial-like amplifying progenitors is enhanced following lesion

Successful neuro-regeneration of brain tissue requires that adult stem cells have the capacity to give rise to newborn neurons over time. To test if pRG and NE-Ap were capable of reparative neurogenesis, we injected zebrafish at 3-dpl with EdU, and provided chase periods ranging from 7-days post injection (dpi) to 8-weeks post injection (wpi; **Fig. 3a, k**, 7-dpi, *n* = 6; 2-wpi, *n* = 5; 4-wpi, *n* = 6; 8-wpi, *n* = 5). Across all chase periods, we were unable to find evidence of newly co-labelled EdU^+^/HuC/D^+^ cells in the PGZ in proximity to the lesion site that could have arisen from the pRG population (**Fig. 3b-c**). Given this result, we asked whether pRG may instead undergo gliogenesis to replace damaged RG following lesion within the qRG layer of the PGZ. In agreement with this hypothesis, we consistently identified a population of co-labelled EdU^+^/GS^+^ cells within the qRG layer following all chase periods (**Fig. 3d-g**). Quantification revealed no significant difference in the EdU^+^ population size throughout the PGZ (**Fig. 3h**; neuronal layer + qRG layer), nor in the newborn EdU^+^/GS^+^ population (**Fig. 3i**) at all time points following EdU injection. By taking the ratio of co-labelled cells to the total EdU^+^ population we found that this cell fraction did not deviate by more than 10% across groups (**Fig. 3j**). These findings propose that pRG are unipotent in the tectum after stab lesion, and function to produce radial-glia, to replenish the qRG cells lost to injury.

**Figure 3.**
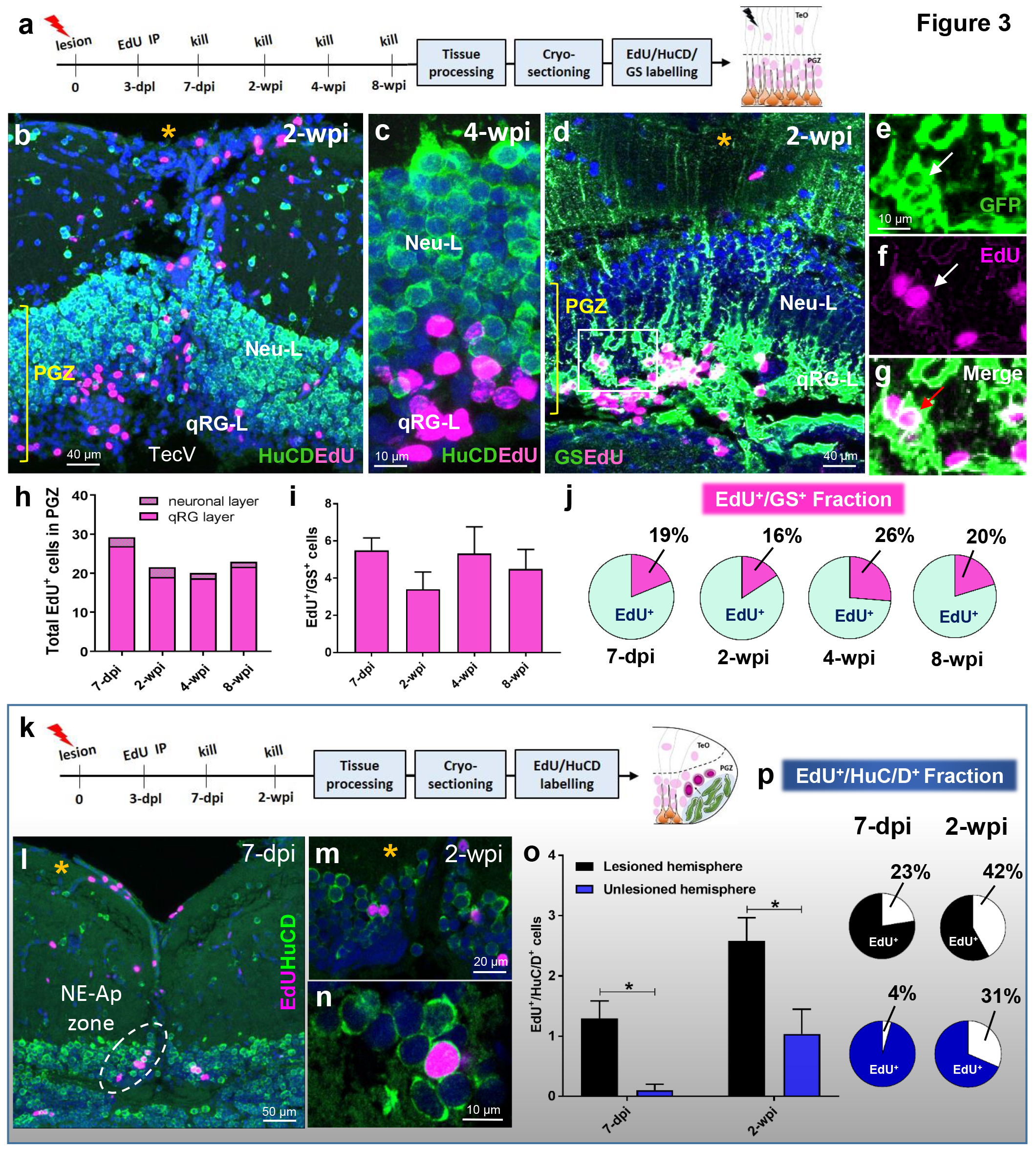
Differentiation post-lesion in stem cell zone 1 (pRG) and zone 2 (NE-Ap). **(a)** Experimental design to study differentiation of proliferating radial-glia (pRG) at chase periods post-EdU injection. **(b-c)** Low and high magnification examples displaying the absence of co-labelling of EdU^+^ cells (pink) in the PGZ with the neuronal marker, HuC/D (green) at 2-wpi (**b**) and 4-wpi (**c**). **(d-g)** Representative images of co-labelling of EdU^+^/GFAP^+^ cells in the qRG layer (qRG-L) of the PGZ at 2-wpi. White box in **d** shown at higher magnification and in separate channels confirming co-labelling of pRG with EdU (**e-g;** white arrows). **(h)** Total EdU^+^ cells in the upper neuronal layer and deeper qRG layer at consecutive chase periods (one-way ANOVA; F (3, 35) = 2.239, *p* = 0.1009; Tukey’s multiple comparisons test: 7-dpi vs 2-wpi, *p* = 0.2082; 7-dpi vs 4-wpi, *p* = 0.1072; 7-dpi vs 8 wpi, *p* = 0.3333; 2-wpi vs 4-wpi, *p* = 0.9863; 2-wpi vs 8-wpi, *p* = 0.9984; 4-wpi vs 8-wpi, *p* = 0.9621). **(i)** Quantification of EdU^+^/GS^+^ cells at increasingly longer chase periods post tectal lesion in the qRG layer (one-way ANOVA; F (3, 17) = 0.7972, *p* = 0.5123; Tukey’s multiple comparisons test: 7-dpi vs 2-wpi, *p* = 0.5190; 7-dpi vs 4-wpi, *p* = 0.9994; 7-dpi vs 8- wpi, *p* = 0.9232; 2-wpi vs 4-wpi, *p* = 0.5845; 2-wpi vs 8-wpi, *p* = 0.9107; 4-wpi vs 8-wpi, *p* = 0.9533). **(j)** Fraction of EdU^+^/GS^+^ cells as a percentage of the total EdU^+^ population in the PGZ at four chase periods examined. **(k)** Experimental design to investigate differentiation of NE-Ap cells to newborn neurons at 7-dpi and 2-wpi. **(l)** Image displaying the midline tectal margin showing the NE-Ap zone of lesioned (yellow asterisk; white hashed lines) and unlesioned hemispheres at 7-dpi. **(m-n)** High magnification examples showing co-labelling of EdU^+^/HuC/D^+^ cells near the tectal margin 2-wpi. **(o)** Number of EdU^+^/HuC/D^+^ cells between the lesioned and unlesioned hemispheres at 7-dpi and 2-wpi of EdU (unpaired t-test, two-tailed: 7-dpi lesion vs 7-dpi unlesioned, *p* = 0.0083; 2-wpi lesion vs 2-wpi unlesioned, *p* = 0.0336). **(p)** Fraction of EdU^+^/HuC/D^+^ cells as a percentage of the total EdU^+^ population in the PGZ across chase times post-lesion. All data presented are mean ± S.E.M. Significance was accepted at \**p* < 0.05.

We next investigated if increased proliferation of NE-Ap after injury resulted in elevated neurogenesis (**Fig. 3k**). Within the NE-Ap zone we identified a significant increase in the co-labelled population of EdU^+^/HuC/D^+^ cells, with more than double the number of EdU^+^/HuC/D^+^ cells by 2-wpi (**Fig. 3l-o**). This observation was further supported by the enlarged fraction of the co-labelled population in the lesioned hemisphere (percentage of white:black in circles) between time points (**Fig. 3p**). These results illustrate that NE-Ap cells at the TMZ of the adult zebrafish increase their constitutive rate of neurogenesis following tectal lesion.

### Physiological levels of Wnt/β-catenin signalling is unaltered in the qRG layer, but increased in the NE-Ap zone following injury

Wnt/β-catenin signalling has repeatedly been shown to be essential for vertebrate midbrain-hindbrain development (Rhinn and Brand, 2001; Buckles et al., 2004; Hüsken and Carl, 2013; Duncan et al., 2015) as well as tissue regeneration within CNS (Ramachandran et al., 2011; Meyers et al., 2012; Wan et al., 2014; Briona et al., 2016) and non-CNS structures of the zebrafish (Stoick-Cooper et al., 2007; Azevedo et al., 2011; Wehner et al., 2014; Stewart et al., 2014;). To investigate the role of Wnt/β-catenin signalling in the cellular response of qRG and NE-Ap populations following tectal lesion, we analysed the expression pattern of downstream markers and target genes of the canonical Wnt pathway under physiological conditions compared with those at 3-dpl (**Fig. 4a**). Using antibodies against β-catenin, we report that baseline levels of nuclear β-catenin within qRG cells is widespread across the qRG layer (**Fig. 4b-d**), but that upon lesion little change in the expression pattern is detected (**Fig. 4e-f**). Repeating these experiments using established transgenic Wnt reporter lines, such as *Tg(TopFlash:gfp)*, a Wnt reporter line containing 4 TCF binding sites upstream of a destabilized GFP transgene (Dorsky et al., 2002), and *Tg(TCFSiam:mCherry)*, a Wnt reporter containing 7 multimerized TCF/LEF binding sites and the *Xenopus* Siamois promoter upstream of mCherry (Moro et al., 2012), recapitulated this same pattern in the qRG layer (**Fig. 4h-i**, control; **Fig. 4j-k**, 3-dpl). Interestingly, we detected a clear increase in the proportion of mature neurons in the superficial PGZ expressing nuclear β-catenin following lesion (**Fig. 4b, e**, yellow arrows), that could additionally be detected within the tectal NE-Ap zone post-injury proximal to EdU^+^ cells (**Fig. 4l-m**; dashed circle). Despite this, no evidence of β-catenin staining within individual NE-Ap cells under control or lesioned conditions was observed. Examination of the downstream Wnt target gene, *axin2*, in the quiescent RG layer and NE-Ap zone confirmed the same pattern of expression between these two stem/progenitor tectal populations (**Fig. 4n-p**, control; **Fig. 4q-s**, 3-dpl). These findings show that while Wnt/β-catenin signalling in qRG following lesion appears unaltered, Wnt signalling is elevated in the NE-Ap zone post-injury.

**Figure 4.**
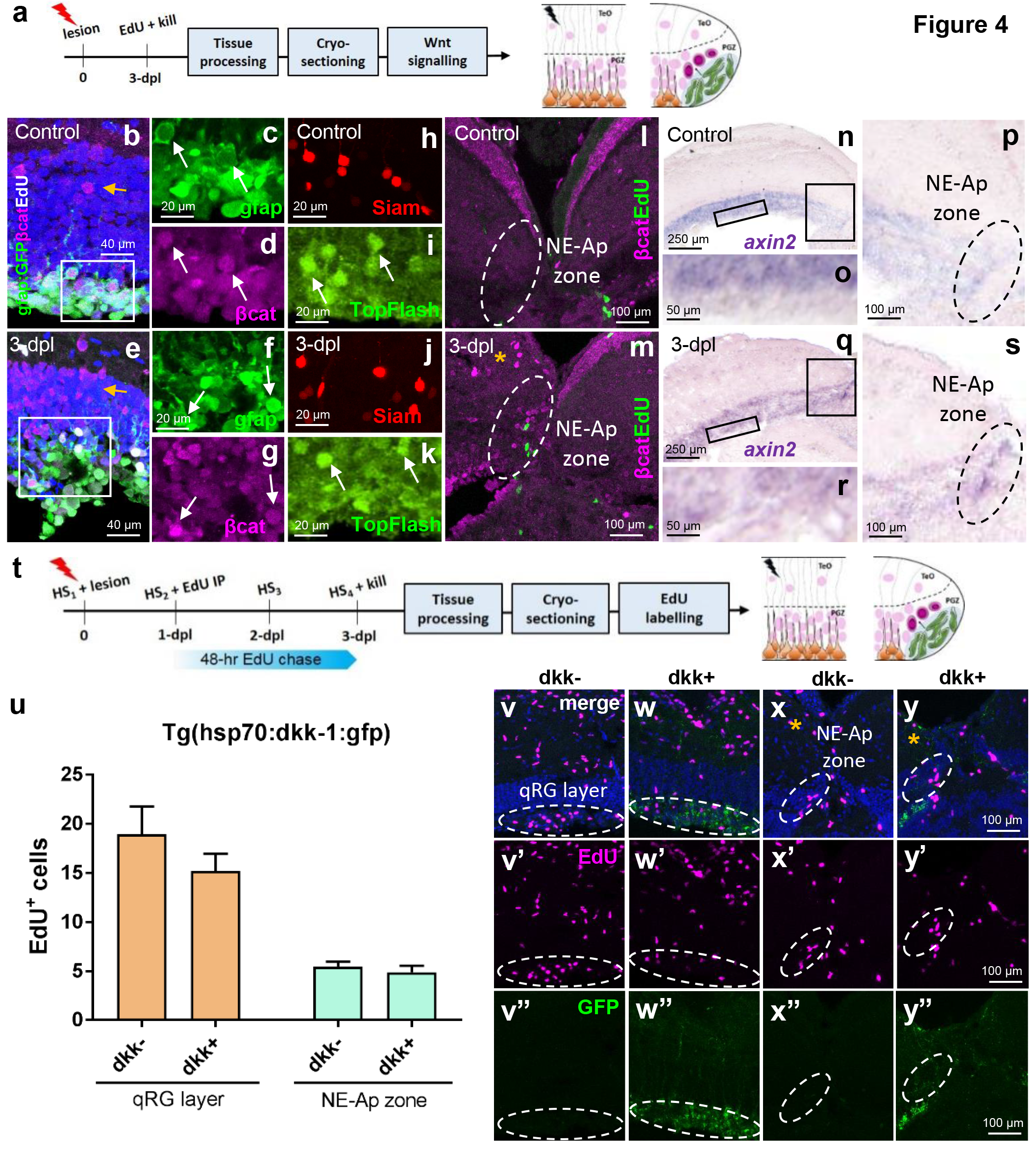
Wnt/β-catenin signalling following tectal lesion in stem cell zone 1 (pRG) and zone 2 (NE-Ap). **(a)** Experimental design to study Wnt/β-catenin signalling at 3-dpl. **(b-d)** Physiological levels of β-catenin expression in the neural layer (Neu-L) and qRG layer (qRG-L; stem cell zone 1). A small number of β-catenin-positive neuronal cell bodies are seen in the Neu-L (yellow arrow). White box depicts higher magnification images of the qRG-L in **c-d**. Split channel images showing co-labelling of qRG (**c**) and β-catenin (**d**; white arrows). **(e-g)** Lesion-induced β-catenin staining in the neural layer (Neu-L) and qRG layer (qRG-L) showing upregulation in the proportion of neuronal cells expressing nuclear β-catenin. White box depicts higher magnification images of the qRG-L in **f-g**. Split channel images showing co-labelling of qRG (**f**) and β-catenin (**g**; white arrow). **(h-k)** Common expression pattern of Wnt observed in the qRG-L between control (**h-i**) and at 3-dpl (**j-k**) using the Wnt reporter lines *Tg(TCFSiam:mCherry)* and *Tg*(*Topflash:gfp*; white arrows). **(l-m)** β-catenin expression and EdU-labelling in the NE-Ap zone (stem cell zone 2; dashed white circles) under control conditions (**l**) and at 3-dpl (**m**, orange asterisk), showing increased expression post-lesion. **(n-p)** Homeostatic levels of *axin2* expression in the qRG layer (**o**) and NE-Ap zone (**p**; black dashed circle). Black boxes in **n** denote higher magnifications in **o-p**. **(q-s)** *axin2* expression 3-dpl in the qRG layer (**r**) and NE-Ap zone (**s**; black dashed circle). Black boxes in **q** denote higher magnifications in **r-s**. **(t)** Experimental design to study the requirement of Wnt/β-catenin signalling for the proliferative response of stem cells to tectal stab lesion using the heat-shock line *Tg(hsp70l:dkk-1:gfp)*. **(u)** EdU population size in the qRG layer and NE-Ap zone shows no change in proliferation post-injury in the absence of Wnt signalling (dkk+) compared with wildtype animals (dkk-) using the *Tg*(*hsp70:dkk-1:gfp*) transgenic line (unpaired t-test, two-tailed: qRG layer, *p* = 0.3081; NE-Ap zone, *p* = 0.4960). **(v-w”)** Representative images of EdU^+^ staining (pink) in the qRG layer (hashed lines) in dkk- (control; **v-v”**) and dkk+ (**w-w”**) post-lesion. **(x-y”)** Representative images of EdU^+^ staining (pink) in the NE-Ap zone (hashed lines) in dkk- (control; **x-x”**) and dkk+ (**y-y”**) post-lesion. Orange asterisk denotes the lesioned hemisphere. Note the GFP-positive expression observed in the dkk+ (**w”, y”**) but not dkk- (**v”, x”**). All data presented are mean ± S.E.M. Significance was accepted at \**p* < 0.05.

### Wnt/β-catenin Signalling is Not Required for Stem/Progenitor Proliferation Post-Injury

To test whether homeostatic levels of Wnt/β-catenin signalling were required for injury-induced proliferation of cells within the qRG layer and NE-Ap zone, we next over-expressed Dkk-1, a negative regulator of Wnt signalling over 4-consecutive days using the *Tg*(*hsp70:dkk-1:gfp*) transgenic line (Stoick-Cooper et al., 2007). We subsequently quantified cell proliferation following a 48-hr EdU chase compared with wildtype levels (dkk-) in the lesioned hemisphere (**Fig. 4t**; wildtype, *n* = 4; dkk+, *n* = 4). Surprisingly, we detected no significant change in the population size of EdU^+^ cells between dkk- and dkk+ fish in either stem cell zone analysed (**Fig. 4u-y**). Taken together, these experiments imply that the proliferative responses within distinct stem/progenitor niches following tectal injury appear to arise independent of Wnt/β-catenin signalling.

## DISCUSSION

Our findings support the notion that niche-specific stem/progenitor cells in the adult zebrafish brain are distinguished by discrete regenerative capacities. We show that both quiescent and active stem/progenitor populations of the adult tectum play an important role in repairing damaged tissue following injury. To date, it has been unclear whether the homogeneous quiescent radial-glial (qRG) population that compose the roof of the midbrain ventricle retain a regenerative program to repopulate lost neurons and glia following tectal injury. Our data highlight that lesion-induced, newly proliferating radial-glia (pRG) of the tectal periventricular grey zone (PGZ) are unipotent producing progeny destined for a radial-glia fate (**Fig. 5** – **top**). Conversely, we report that constitutively active neuro-epithelial-like amplifying progenitors (NE-Ap) of the tectal marginal zone (TMZ) that contribute to lifelong neurogenesis, additionally respond to injury by increasing their normal rates of proliferation and neurogenesis alongside elevated levels of Wnt/β-catanin signalling (**Fig. 5**-**bottom**). The balance by which quiescent versus constitutively active stem/progenitor cells respond to neuro-trauma in the CNS remains poorly understood. Such diversity in the injury response of niche-specific stem and progenitor cells has been documented in the hippocampal dentate gyrus (Jhaveri et al., 2015; Gao et al., 2009) and subventricular zone (SVZ) of the adult mammalian brain (Thomsen et al., 2014). The results of our study reflect our earlier work examining neurogenic plasticity in stem cell compartments of the adult zebrafish brain, where we reported that upon changes to the social environment (Lindsey and Tropepe, 2014) or sensory stimuli (Lindsey et al., 2014), NE cells appear more intrinsically primed to modulate their cellular behaviour compared with RG. Understanding the molecular differences between the newly identified, slowly-cycling *Her5*-positive NE stem cells in the caudal tectum (Alunni et al., 2010; Galant et al., 2016; Dambroise et al., 2017) and qRG of the midbrain tectum, along with the actively cycling RG in the telencephalon would further aid to determine how quiescence and proliferative activity is controlled.

**Figure 5.**
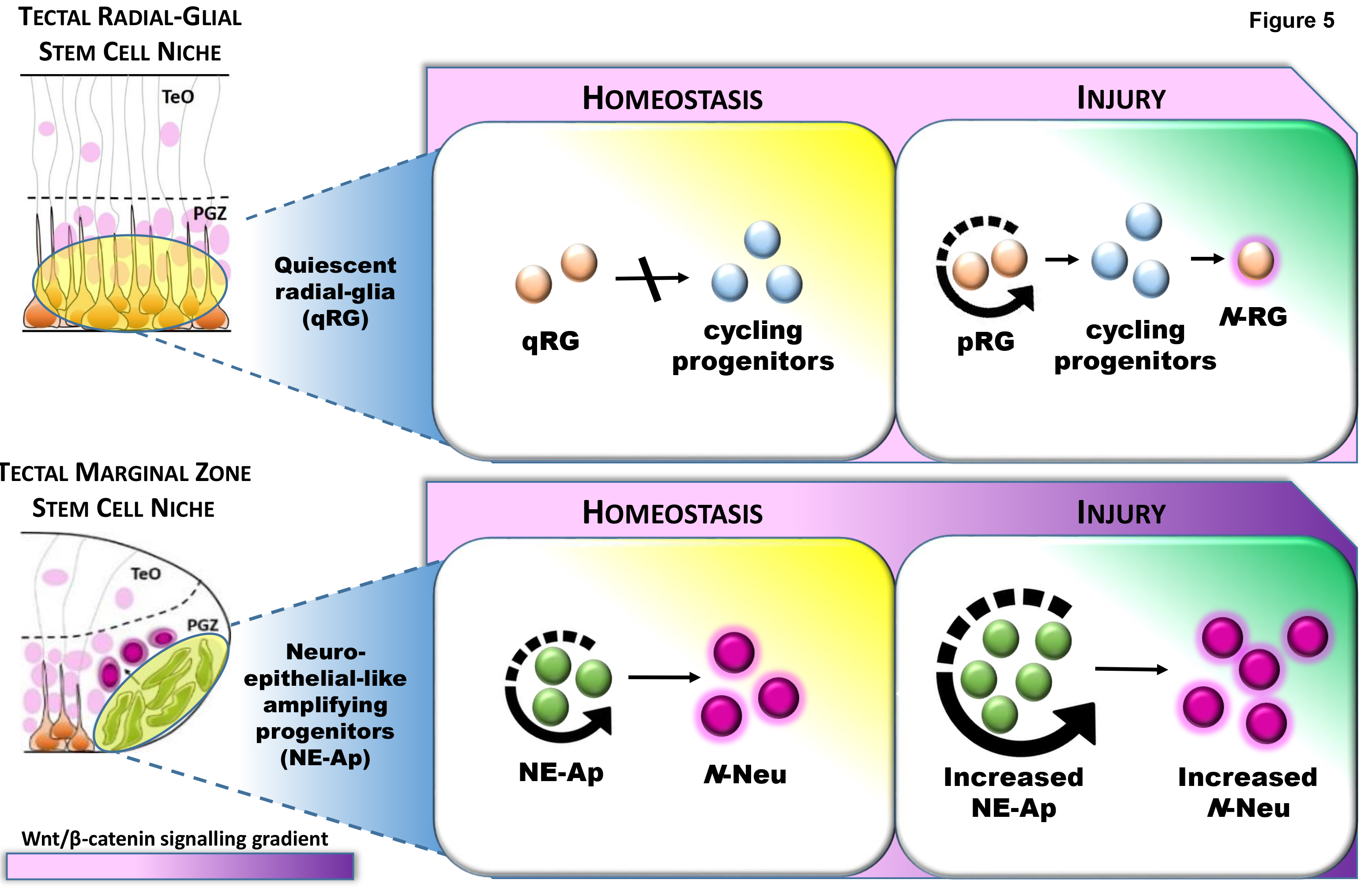
Model displaying the response of niche-specific stem/progenitor populations following lesion to the adult optic tectum. **Top:** Quiescent radial-glia (qRG) are unable to produce cycling progenitors under homeostasis at the tectal ventricle. Post-injury, a subpopulation of qRG transition to a proliferative state (pRG) to produce newly cycling progenitors which differentiate into *de novo* radial-glia (*N-*RG). **Bottom:** Neuro-epithelial-like amplifying progenitors (NE-Ap) are constitutively active under homeostasis, acting as a life-long source of newborn neurons (*N*-Neu) at the tectal marginal zone. Tectal injury induces NE-Ap to increase their homeostatic level of cell proliferation, leading to an increase in neuronal output to replace neurons lost to damage. This regenerative process is associated with an increase in Wnt/β-catenin signalling. TeO, tectum opticum; PGZ, periventricular grey zone.

Vertebrate neural stem cell niches are characterized by heterogeneity in the stem/progenitor population (Shen et al., 2006; Merkle et al., 2007; Ganz et al., 2010; Chaker et al., 2016; Magnusson and Frisén, 2016). Accumulating evidence shows that such heterogeneity also may exist amongst quiescent stem and progenitor cell populations. Single-cell analyses of quiescent neural progenitors in both the SVZ and hippocampus show that subpopulations within a niche can be distinguished by diverse characteristics such as their response to hormones, brain ischemia, metabolic state, and growth factors (Jhaveri et al., 2015; Llorens-Bobadilla et al., 2015; Shin et al., 2015; Luo et al., 2015). This indicates that different quiescent precursors are likely governed by varying degrees of plasticity. Morphological analysis has also revealed that a subpopulation of quiescent B1 cells of the adult ventricular-SVZ are defined by the presence of envelop-limited chromatin sheets, uniquely discriminating this population within the forebrain neurogenic niche (Cebrián-Silla et al., 2017). Across adult zebrafish neurogenic zones, niches of the forebrain dorsal telencephalon and hindbrain vagal lobe are exemplary of domains dominated by heterogeneous RG stem/progenitor populations (Lindsey et al., 2012; Lindsey et al., 2014). The majority of the RG population in these niches reside in a non-cycling state under homeostasis, with only a small number of cells taking up proliferative markers with short pulse-chase experiments. With injury to the forebrain however, a significant proportion of qRG re-enter the cell cycle (Kroehne et al., 2011; Buamgart et al., 2011; Marz et al., 2011; Kishimoto et al., 2012; Barbosa et al., 2015). Impressively, these activated RG in the telencephalon appear to possess the molecular code to repopulate the full complement of lost neuronal lineages (Kroehne et al., 2011).

The RG tectal stem cell niche composes the entire roof of the midbrain ventricle and displays lifelong quiescence apart from RG that are thought to act as amplifying progenitors in proximity to NE cells of the TMZ (Grandel et al., 2006; Ito et al., 2010; Galant et al., 2016). The tectal population of qRG therefore contrasts the mixture of proliferative and non-proliferative RG observed in the forebrain and hindbrain. In this study we proposed that the qRG population may be endowed with the intrinsic program for neuroregeneration, however our evidence points towards this population as serving a more structural role in maintaining the epithelial barrier and supporting the tectal circuitry. In agreement, our data shows limited production of newborn RG progeny from pRG (**Fig. 3i**), along with the lack of neurons produced locally to replace those damaged in upper neuronal layer of the PGZ. This demonstrates that qRG contribute principally to maintaining the structural integrity of the ventricular epithelium towards the tectal ventricle. Functionally, this raises the question of whether neurons adjacent to the stab lesion canal are sufficient for ongoing information processing or if animals maintain a sensory deficit until the columns of newly produced neurons from TMZ NE-Ap cells are shifted laterally to repopulate the lesion site.

A particularly interesting finding of our study was that NE-Ap cells underwent elevated rates of constitute proliferation and neurogenesis following lesion to the midbrain nearly ~350 μm from the TMZ. In the uninjured context, NE-Ap function to circumferentially add newborn neurons to the tectum to maintain tissue growth (Ito et al., 2010). Our data indicate that signals initiated at the lesion site were received by NE-Ap of the lesioned hemisphere leading to increases in both cell proliferation (**Fig. 2p**) and neurogenesis (**Fig. 3p**)
beyond changes detected in the unlesioned hemisphere. This finding argues the regenerative mode of the tectal NE-Ap population and whether it may be viewed as compensatory neurogenesis, rather than lineage-directed neural regeneration. Compensatory neurogenesis following CNS damage generally repairs and strengthens the circuitry without the necessity to directly replace lost cells and could therefore be seen as a broader restorative mechanism (Kaslin et al., 2008). In line with our data, as well as previous work in the telencephalon (Kroehne et al., 2011), increases above constitutive levels of cell proliferation and/or neurogenesis normally produced from a stem cell niche have also been associated with a compensatory response. Mammalian studies of disease and injury have commonly shown the occurrence of compensatory proliferation/neurogenesis as an endogenous response to CNS tissue repair (reviewed in Goldman et al., 2005). Stab lesions immediately within the zebrafish TMZ would be a fruitful approach to examine the potential of NE-Ap for regenerative neurogenesis and gliogenesis, and to conclude the specific regenerative mode employed by NE-Ap cells.

A common form of neuroregeneration brought to light by comparing both the adult cerebellar and tectal niches, is the discovery that NE, rather than RG, appear to drive neuroregeneration in the mid and hindbrain. RG have often been proposed as the major stem cell population regulating regeneration in vertebrates, however here we provide additional evidence supporting the notion that this is likely not the case. In the injured cerebellum, NE are able to produce all cell lineages that they normally produce under homeostasis (Kaslin et al., 2017). In agreement, we show for the first time that, in the tectum, neuroregeneration from a constitutively active population of NE-Ap functions similarly to populate the neural lineage that is not replenished by pRG cells. Therefore, in line with NE-driven cerebellar regeneration, midbrain NE-Ap also make use of their homeostatic neurogenic program.

Amongst the cues controlling the stem cell state, developmental signalling pathways play an important role (Ganz et al., 2010; Chapouton et al., 2010; Rothenaigner et al., 2011; Alunni et al., 2013; de Oliveira-Carlos et al., 2013; Rodgers et al., 2014; Wang et al., 2012; Daynac et al., 2016; Kawai et al., 2017). Here, we investigated the role of Wnt/β-catenin signalling in regulating qRG and NE-Ap during the regenerative response given the importance of this pathway for early midbrain-hindbrain boundary patterning (Buckles et al., 2004; Duncan et al., 2015) and tissue-wide regeneration in the zebrafish (Stoick-Cooper et al., 2007; Ramachandran et al., 2011; Azevedo et al., 2011; Meyers et al., 2012; Wan et al., 2014; Stewart et al., 2014; Duncan et al., 2015; Briona et al., 2016). It has recently been found that Wnt signalling components play a role in constitutive cell division and neuronal differentiation from NE-Ap. Inhibitors of Wnt, such as IWR-1, further suppress proliferation of this progenitor population under homeostasis (Shitasako et al., 2017). Using a combination of downstream Wnt targets, including β-catenin, TCF/Lef, Siam, and *axin2*, we show a baseline level of canonical Wnt signalling in both qRG cells of the tectum and within the TMZ (**Fig. 4**). We find however, that upon tectal lesion Wnt/β-catenin signalling is upregulated uniquely in the TMZ where NE-Ap reside (**Fig. 4m, s**). This finding led us to hypothesize whether the increase in cell proliferation from the NE-Ap population with injury was controlled by the Wnt/β-catenin pathway. Silencing Wnt signalling over the first 3-days post-lesion by overexpression of the Wnt inhibitor, *Dickkopf-1*, displayed no significant difference in the cycling population however (**Fig. 4u-y**). This suggests that despite elevated levels of Wnt/β-catenin signalling following injury, Wnt may not be required for the regenerative response of NE-Ap or may be acting in combination with other signalling cascades as illustrated in the zebrafish retina (Wan et al., 2014). Our results align with studies of the larval zebrafish hypothalamus showing that partial genetic ablation of hypothalamic RG leads to a net increase in cell proliferation in the absence of Wnt/β-catenin signalling (Duncan et al., 2015).

Earlier studies of the zebrafish CNS have highlighted the heterogeneous role of Wnt/β-catenin signalling in controlling stem cell behaviour. A pointed example of this is in the zebrafish retina. The retina contains populations of ciliary marginal zone (CMZ) NE cells and quiescent Müller glia reflecting the region-specific organization of the tectal marginal zone (TMZ) NE-Ap and qRG. In this model, Wnt/β-catenin signalling is required to permit normal retinal growth from the CMZ, whereas activated Müller glia depend on these signals to generate neuronal progenitors post-lesion (Meyers et al., 2012). The zebrafish spinal cord instead depends on the Wnt pathway for both RG-derived neurogenesis and axonal regrowth post transection (Briona et al., 2016; Birona and Dorsky, 2014). Additionally, Notch signalling regulates stem cell quiescence (Chapouton et al., 2010; Rothenaigner et al., 2011; Alunni et al., 2013) in distinct stem/progenitor populations. Therefore, uncovering how Wnt/β-catenin signalling may associate with Notch signalling in diverse physiological and pathological states would serve well to resolve the contribution of these two heavily studied pathways to developmental and regenerative processes in the zebrafish CNS.

In the present study we have illustrated that tectal stab lesion leads to a dichotomous response in cell proliferation and differentiation between activated qRG and NE-Ap. Signals produced as a result of injury are sufficient for these stem and progenitor populations to initiate their individual reparative programs to replace damaged RG and neurons. While the relative contribution and mode of CNS regeneration differs between these adult stem/progenitor populations, our results propose that they complementarily possess the cellular and molecular machinery needed to restore tissue back to its uninjured state over time. Our findings are important in contributing new evidence that the regenerative behaviour of diverse adult stem/progenitor phenotypes contain different intrinsic degrees of regenerative plasticity and regulation in the context of injury. Moving forward, defining the transcriptional profiles of closely related stem/progenitor phenotypes residing in different neurogenic domains under homeostasis and with injury will be fundamental in mapping their distinct molecular programs.

## METHODS

### Animals

All zebrafish lines used in experiments were housed in the Monash University FishCore Facility and maintained according to facility regulations. Animals across experiments were of mixed sex and between 6-months to 1-year of age. For individual experiments animals were age and size matched. Upon completion of experiments, zebrafish were sacrificed using an overdose of 0.4% Tricaine (Sigma; E10521) diluted in ice-cold facility water. Experiments were assessed and approved by the Monash University Animal Ethics Committee and were conducted under applicable Australian laws governing the care and use of animals for scientific research.

### Optic Tectum Stab Lesion Assay

Adult fish were anesthetized in 0.04% Tricaine (Sigma; E10521) prior to tectal stab lesion. Stab lesions were performed under a dissecting microscope in the left hemisphere of the tectum using a 30 × ½ gauge canula (Terumo). The cannula was carefully inserted vertically into the centre of the neurocranium overlying the tectum, and slightly anterior to the middle of the tectum (**Fig. 1g-h**). These landmarks ensured the lesion entered through the tectal layers and into the large, underlying tectal ventricle. Thereafter, fish were returned to experimental tanks and monitored until normal swimming behaviour commenced. Sham fish (controls) were treated similarly, with the exception that the cannula only penetrated the neurocranium, but not further into the brain tissue.

While we found our tectal lesion assay to be highly reproducible, to ensure accurate cell quantification of the lesion response across experiments the following screening criteria had to be met in order for the brain sample to be included in downstream analyses: (1) lesioned hemispheres displayed a clear upregulation of the proliferative marker (EdU or PCNA), and (2) the cannula clearly penetrated the deep PGZ layer as per 4,6-diamidino-2-phenylindole (DAPI) counterstaining.

### EdU Injections

A 10 mM solution of 5-ethynyl-2’-deoxyuridine (EdU; ThermoFisher; A10044) diluted in 1X-Phosphate Buffered Saline (PBS; pH 7.4) was injected intraperitoneally to label proliferating cells in the *S*-phase of the cell cycle. The total volume of individual EdU injections was 40 μL.

### Tissue Processing for Immunohistochemistry, Histology, and *in situ* Hybridization

Following sacrifice, brains were exposed by removing the neurocranium and left *in situ* in the head. Zebrafish were subsequently decapitated, and heads then immediately fixed in 2% paraformaldehyde (PFA; Sigma; 158127) diluted in phosphate buffer (pH 7.4) for immunohistochemistry (IHC) or in 4% PFA diluted in 1X-PBS (pH 7.4) for *in situ* hybridization (ISH) and histology, overnight at 4 °C. The next day, heads were placed in a solution of 8 M ethylenediaminetetraacetic acid (EDTA) for 24-hrs at 4 °C to decalcify surrounding cartilage, before being cryo-embedded in a mixture of fish gelatin (Sigma; G7041) and sucrose (VWR; VWRC0335) in PBS. Heads were sectioned in the cross-sectional plane using a Leica CS3080 Cryostat at a thickness of 18 μm and stored at −80 °C until use.

### Tissue Processing and Whole Brain EdU Staining for Optical Projection Tomography

Tissue processing and whole brain EdU staining on adult wildtype (Tübingen) zebrafish for Optical Projection Tomography (OPT) experiments was completed as described previously (Lindsey et al., 2018; Lindsey and Kaslin, 2017). Briefly, brains were excised from the neurocranium and immediately fixed in ice-cold 2% PFA overnight. The next day, brains were rinsed over 4-hrs in 1X-PBS containing 0.3% Triton X-100 (Tx; Sigma; T9284) to remove fixative, then placed in a solution of 1% Tx/5% dimethyl sulfoxide (DMSO; Millipore; 317275) in 1X-PBS for 24-hrs to increase antibody penetration. Thereafter, brains were incubated in EdU staining mixture for 4-days with gentle agitation, then rinsed before embedding in low melting agarose (Sigma; A9414). Trimmed agarose blocks were dehydrated in pure methanol over 1.5-days, and finally cleared in a 2:1 solution of benzyl benzoate:benzyl alcohol (BABB; Sigma; B6630, 402834).

### EdU Staining on Cryosections

Rehydrated cryosections were stained as per the “Click-iT” reaction between EdU and an azide-modified Alexa dye (Molecular Probes, 2010). Slides were incubated at room temperature in the dark for 30-min in a cocktail consisting of 1X-PBS, 0.5M L-Ascorbic Acid (Sigma; A5960), 2M Tris buffer (pH 8.5; Sigma; T6791), Copper II Sulphate (Sigma; C1297), and 100mM Azide Fluor dissolved in DMSO (555 or 647; ThermoFisher; A20012; A10277). A total volume of 250 μL of staining mixture was used per slide. Counterstaining was performed using DAPI.

### Hematoxylin & Eosin Histology

Hematoxylin & eosin (H&E) histological staining was completed at the Monash Histology Platform at Monash University using a Leica ST5010 Autostainer and CV5030 Coverslipper. Imaging was completed using an Olympus Provis AX70 Widefield brightfield microscope at the Monash Micro Imaging (MMI) facility at Monash University.

### Immunohistochemistry

Immunohistochemistry was performed as previously described (Grandel et al., 2006). Briefly, rehydrated cryosections were incubated in primary antibody overnight at 4 °C. Tissue was then incubated with secondary antibodies (dilution 1:750) conjugated to Alexa Fluor 488, 555, and 633 (Invitrogen) for 1-hr at room temperature and counterstained with DAPI. For confocal imaging, all sections were mounted in 70% glycerol in 1X-PBS and cover-slipped.

For HuC/D staining, antigen retrieval was completed by incubating sections in 50 mM Tris buffer (pH = 8.0) for 30-min at 80-85 °C in a tissue incubator. Thereafter, tissue was stained as described previously (Lindsey et al., 2012). For labelling of Proliferating Cell Nuclear Antigen (PCNA), tissue was incubated in 10 mM sodium citrate buffer (pH 6.0) for 15-min at 90 °C.

Primary antibodies used included mouse monoclonal against glutamine synthetase (GS, 1:4000; Millipore; MAB302), rabbit polyclonal against S100β (1:1500; Dako; Z0311), mouse monoclonal against PCNA (PC10, 1:750; Dako; M0879), mouse monoclonal against β-catenin (15B8, 1:1500; Santa Cruz; sc-53483), mouse polyclonal against HuC/D (1:400; Molecular Probes; 16A11), and rabbit polyclonal against Green Fluorescent Protein (GFP, 1:1000; ThermoFisher; A11122).

### Transgenic Lines

The *Tg*(*gfap:GFP*)*^mir2001^* line was used to label the quiescent radial-glial (RG) population at the tectal ventricle. *Tg*(*mpeg1:mCherry*)*^g122^* was used to label resident microglia and infiltrating macrophages upon lesion, and was a kind gift from Dr. G. Lieschke at the Australian Regenerative Medicine Institute (ARMI). *Tg*(*Topflash:gfp* (Dorsky et al., 2002) and *Tg*(*TFCsiam:mCherry* (Moro et al., 2012) were used to assess canonical Wnt expression. The heat-shock inducible *Tg*(*hsp70:dkk-1:gfp* (Stoick-Cooper et al., 2007) line was applied to negatively regulate Wnt signalling via *Dickkopf-1* (Dkk-1) with heat-shocks performed as described below.

### *In Situ* Hybridization

*In situ* hybridization on cryosections from adult wildtype zebrafish (Tübingen) was performed as described previously (Ganz et al., 2014). The *axin2 in situ* probe was generated in our lab from the ZP60 Axin2 plasmid (Addgene #16882) in a pSPORT1 vector, digested with asp718 and transcribed with SP6 RNA polymerase. The probe was used at a concentration of 1:250.

### Heat-Shock Treatments

Adult *Tg*(*hsp70:dkk-1:gfp*) fish were heat-shocked once daily over four consecutive days to reveal GFP expression and downregulate Wnt/β-catenin signalling. Daily heat-shock treatments were performed in the morning by placing fish into the heat-shock chamber with the water at an initial temperature of 28 °C and the heater turned on to increase the temperature to 38.5 °C over the next hour. Fish were then maintained in the chamber at 38.5 °C for 30-min, before the heater was switched off and the water was progressively returned to 28 °C throughout the remainder of the day. During all heat-shock trials fish were monitored to ensure health was not compromised. Animals negative for the dkk-1 transgene were used as controls and exposed to the same experimental protocol.

### Neuroanatomical Regions of Interest for Imaging and Analysis

#### Periventricular Grey Zone (PGZ) of the optic tectum

The PGZ of the optic tectum, consisting of the upper neuronal layer and deeper quiescent radial-glial layer, was examined between anterior-posterior levels 160-170 as per the adult zebrafish atlas (Wullimann et al., 1996; **Fig. 1f**; ***stem cell zone 1***). Tissue was investigated specifically at the level in which the lesion occurred, unless stated otherwise. This neuroanatomical domain permitted assessment of changes in the proliferative activity and Wnt expression of qRG populations between lesioned and unlesioned conditions in addition to changes in Wnt expression in neuronal layers.

#### Tectal marginal zone

The tectal marginal zone (TMZ) was examined at the same anterior-posterior level as the PGZ to analyze neuro-epithelial-like amplifying progenitor (NE-Ap) proliferation, neurogenesis, and Wnt expression between lesioned and unlesioned conditions (**Fig. 1f**; ***stem cell zone 2***).

#### Caudal tectum

The caudal tectum, consisting of a strip of cells, spanning anterior-posterior levels 208-213 (Wullimann et al., 1996) was investigated to assess changes in EdU proliferation in the caudal NE progenitor (NE-p) domain (**Fig. 1f; *stem cell zone 3***), where defined populations of NE parent stem cells, label-retaining NE progenitor cells, and transit amplifying NE progenitors reside (Galant et al., 2016).

### Optical Projection Tomography (OPT) Imaging and Analysis

EdU-stained adult zebrafish brains were affixed to metal OPT mounts and imaged using a Bioptonics 3001 OPT scanner (Bioptonics, Edinburgh, UK). For each channel (555nm; 488nm) 2-dimensional images were acquired at 0.45° intervals, with each frame averaged twice. EdU-labelling was imaged in the 555nm channel, while brightfield images of brain volume were captured by imaging auto-fluorescence from the 488nm channel. Exposure was consistently set at 75% of the maximum signal intensity detected in the brains. 2-dimensional images were then post-processed using nRECON software (Bruker microCT) to yield a final 3-dimensional dataset for visualization and downstream volumetric analysis of changes in EdU using IMARIS software.

To assess the proliferative response within the lesioned hemisphere following stab injury compared with uninjured conditions, we designed an algorithm in IMARIS to calculate the total volume of EdU^+^ labelling in concentric spherical rings radiating outwards from a centre-point (**Fig. S1a-b**). Here, a centre-point was placed at the junction of the tectal ventricle and qRG-layer where we normally quantify the lesion response (**Fig. S1c – yellow dot**). From this centre-point, we generated concentric spheres at 50 μm intervals, terminating at 299 μm from the centre-point (i.e. 6 spheres totals; **Fig. 1d-l**). Our algorithm then calculated the total EdU volume occupied within each consecutive sphere. We predicted that compared to control levels where nearly no cycling cells are observed, following lesion an increase in the total amount of EdU should be detected in one or more spheres. For example, the total EdU volume within the blue sphere (**Fig. S1f**) ranging from 100-149 μm from the centre-point (**Fig. S1c**) could be calculated independent of other spheres located more proximal or distal to the lesion site. For normalization, the ratio of EdU volume: sphere volume was calculated, and converted to a proportion (represented as percentage) for statistical analysis and graphical representation.

### Confocal Image Acquisition and Processing

Confocal imaging was completed using a Leica SP5 inverted channel confocal microscope with 20X, 40X, 63X, and 100X oil immersion lenses at the Monash Micro Imaging facility (MMI, Monash University). Virtual zoom was applied in cases where cellular detail or co-localization was required. Images shown are maximum projections of z-stacks taken at 1 μm intervals through the z-plane of cryosectioned tissue at 1024^2^ resolution. In all images, dorsal is oriented upwards. For display, brightness and contrast were adjusted using FIJI/ImageJ or LAS AF Lite software.

### Cell Quantification

For all experiments, 2-3 sections at the neuroanatomical region of interest (**Fig. 1f**; stem cell zones 1-3) in each brain sample were analysed. Quantifications were done by counting cells through the z-stack of confocal images taken at 40X-63X magnification using FIJI/ImageJ or LAS AF Lite software to avoid double counting. For co-labelling experiments, orthogonal views were used to accurately confirm double-positive cells. For each brain section, a sum of the cell population of interest was obtained. Sums from multiple brain sections of a single brain sample were then averaged, with this value representing the average number of cells counted for a single biological specimen.

### Statistical Analysis and Graphical Representation

GraphPad Prism7 was used for all statistical analyses and graphs. Statistics were performed on the average cell counts/section calculated across multiple brain samples. For all experiments, significance was accepted at *p* < 0.05. The Shapiro-Wilk normality test was used to confirm data adhered to a Gaussian distribution. Unpaired t-tests (two-tailed) were used to compare differences between two groups. Comparisons between greater than two treatment groups were completed using one-way ANOVA. For all one-way ANOVA tests, the F-value and exact *p*-values are stated. In cases where a significant between-group effect was present, Tukey’s multiple comparisons tests were performed and the adjusted *p*-value reported. All statistical results, including exact p-values, are reported in figure legends. Graphical data represent mean ± standard error of the mean (S.E.M.). Fractions shown depict the total number of co-labelled cells divided by the total EdU^+^ population size, and are represented in percentage.

## ACKNOWLEDGEMENTS

We thank the Monash Micro Imaging (MMI) Facility and the zebrafish FishCore for excellent support. Specific thanks to Dr. K. Schulze, Image Analysist specialist at MMI for his assistance in developing the algorithm to analyse our Optical Projection Tomography (OPT) data. We thank the laboratory of Prof. I. Smyth in the Department of Anatomy and Developmental Biology at Monash University for the continued use of their Optical Projection Tomography scanner for 3-dimensional analysis of our tectal lesion assay. Kind thanks to Prof. G. Lieschke for providing the *Tg*(*mpeg1:mCherry*)*^g122^* macrophage line. Thank you to A. Douek for helpful comments in improving this manuscript. This work was supported by an NHMRC project grant (GNT1068411), Monash University Faculty of Medicine and Nursing strategic grant and Operational Infrastructure Support from the Victorian Government. BWL was supported by a fellowship from NSERC in Canada.

## AUTHORS CONTRIBUTIONS

J.K. and B.L. conceived experiments. B.L. and G.A. conducted all experiments for the study. J.T., C.V., and M.K. assisted with tissue processing, immunohistochemistry, and confocal imaging. B.L. analysed all data sets. B.L. drafted all figures and manuscript with conceptual input from J.K. All authors reviewed and edited the manuscript for final submission.

---

**Supplementary Figure 1.** Sphere analysis of cell proliferative following tectal lesion. **(a-b)** Mid-horizontal (**a**) and mid-sagittal (**b**) views of a reconstructed adult brain denoting the sequential, coloured, concentric spherical rings spaced 50 μm apart for analysis of the total EdU volume/sphere volume. **(c)** Image of the centre-point of analysis (yellow circle) positioned at the junction of the quiescent RG layer and the tectal ventricle, overlayed on a control brain (EdU, pink). **(d-i)** Example of increasing sphere distance from centre-point (***see* c**) used for analysis and overlayed on a control brain (EdU, pink). **(j)** Experimental design to investigate the proliferative response using EdU labelling in the lesioned tectal hemisphere at 1-dpl and 3-dpl. **(k)** Proportion of EdU volume/sphere represented as a percentage between consecutive spheres at 1-dpl. Proliferation was greatest at 1-dpl, displaying a progressive reduction in the proportion of EdU volume/sphere towards spheres more distal to the centre-point (one-way ANOVA: F (5, 54) = 1.311, *p* = 0.2731). **(l1-l7)** Reconstructed total volume of EdU occupying consecutive spheres at 1-dpl shown *in situ* across all spheres (**l1**, 0-299 μm from centre-point), and additively in overlayed spheres (**l2-l7**). **(m)** Proportion of EdU volume/sphere represented as a percentage between consecutive spheres at 3-dpl. A significant decrease was observed between the proportion of EdU nearest the centre-point (i.e. 0-49 μm) compared with more distal spheres (one-way ANOVA: F (5, 35) = 3.463, p = 0.0120; Tukey’s multiple comparisons test: 0-49 *μ*m vs 50-99 μm, *p* = 0.0294; 0-49 μm vs 100-149 μm, *p* = 0.0421; 0-49 μm vs 150-199 μm, *p* = 0.0441; 0-49 μm vs 200-249 μm, *p* = 0.0307; 0 – 49 μm vs 250-299 μm, *p* = 0.0647). **(n1-n7)** Reconstructed total volume of EdU occupying consecutive spheres at 3-dpl shown in situ across all spheres (**n1**, 0-299 μm from centre-point), and additively in overlayed spheres (**n2-n7**). All data presented are mean ± S.E.M. Significance was accepted at \**p* < 0.05.

**Supplementary Figure 2.** The proliferative response of the adult brain to injury is conserved independently of the site of lesion. **(a-b)** Side-by-side comparison of the proliferative response of the adult brain to telencephalic (EdU, green) and tectal (EdU, pink) stab lesions using OPT imaging to visualize the proliferative pattern across the neuro-axis under homeostasis (i.e. control) and at 1-, 3, and 7-dpl. Note the global upregulation in EdU-labelling at 1-dpl compared with more localized cell proliferation at the injury site at 3-dpl. All brains are shown in dorsal view. White dashed circles denote the brain hemisphere in which the lesion was performed. Dpl; days post lesion.

**Supplementary Figure 3.** Macrophage response to tectal lesion. **(a)** Experimental design to investigate macrophage recruitment and proliferation at the site of lesion at 1-dpl and 3-dpl using the *Tg*(*mpeg1:mCherry*)*^g122^* reporter line. **(b-d)** Distribution of resident microglia under homeostasis in the PGZ and superficial tectal layers (**b**) compared to increase in peripheral and local macrophages at 1-dpl (**c**) and 3-dpl (**d**) at the lesion canal. Red circles denote a change in macrophage morphology from an enlarged cell body with ramified processes under uninjured conditions (**b**) to a condensed, amoeboid-like state post-injury (**c**). **(e-g)** Co-labelling with EdU demonstrated an increase in EdU-labeling post-injury (pink, **e** vs. **f-g**) near the lesion canal, and a small number of proliferating macrophages EdU^+^/mCherry^+^ at 1-dpl and 3-dpl (**f-g**, white arrows). In **c-d**, **f-g**, orange asterisk depicts the site of lesion canal.

